# Rigidified Scaffolds for 3 Angstrom Resolution Cryo-EM of Small Therapeutic Protein Targets

**DOI:** 10.1101/2022.09.18.508009

**Authors:** Roger Castells-Graells, Kyle Meador, Mark A. Arbing, Michael R. Sawaya, Morgan Gee, Duilio Cascio, Emma Gleave, Judit É. Debreczeni, Jason Breed, Chris Phillips, Todd O. Yeates

## Abstract

Numerous technical advances have made cryo-EM an attractive method for atomic structure determination. Cryo-EM is ideally suited for large macromolecular structures, while problems of low signal-to-noise prevent routine structure determination of proteins smaller than about 50 kDa. This size limitation excludes large numbers of important cellular proteins from structural characterization by this powerful technique, including many cell-signaling proteins of high therapeutic interest. In the present work, we use molecular engineering techniques to rigidify an imaging scaffold, based on a designed protein cage, to the point where 3 Å resolution can be achieved, even for very small proteins. After optimizing the design of the rigidified scaffold on test proteins, we apply this imaging system to the key oncogenic signaling protein KRAS, which represents an outstanding challenge in the area of structure-based drug design. Despite its 19 kDa size, we show that the structure of KRAS, in multiple mutant forms, and bound to its GDP ligand, can be readily interpreted at a resolution slightly better than 3.0 Å. This advance further expands the capability of cryo-EM to become an essentially universal method for protein structure determination, including for applications to small therapeutic protein targets.

## Introduction

Cryo-electron microscopy (cryo-EM) is a rapidly expanding method for determining the atomic structures of large molecular assemblies. It is, however, problematic for determining the structures of small-to-medium sized protein molecules. A size of about 38 kDa represents a likely theoretical lower limit (Henderson, 1995), while about 50 kDa is a practical limit from current work (Herzik *et al*. 2019). For comparison, the average eukaryotic protein chain is about 35 kDa in mass, while bacterial proteins are even smaller on average (Yeates *et al*. 2020). Accordingly, vast numbers of important protein targets remain out of reach of this expanding structural method.

The idea of binding a small protein of interest (the ‘cargo’) to a much larger carrier (the ‘scaffold’) in order to make it large enough to visualize by cryo-EM goes back many years (Coscia *et al*., 2016; Kratz *et al*., 1999; Martin *et al*., 2016). A key challenge is how to make the binding attachment between the scaffold and the cargo protein sufficiently rigid, as even minor flexibility in the attachment severely compromises the ability to reconstruct a high-resolution image of the bound cargo component. In addition, a general solution to the scaffolding problem calls for modular design, *i.e*. through the use of a scaffolding component that can be readily diversified to bind any given cargo protein of interest. Among proteins in use as modular binding components, two types under particularly active development include: nanobody and other antibody variants (*e.g*. McMahon *et al*., 2018, Morison *et al*., 2021), and DARPins (*e.g*. Binz *et al*., 2004, Rothenberger *et al*., 2022); DARPins are engineered proteins, structurally distinct from antibodies but analogous by having sequence-variable loop regions that can be selected to bind diverse target proteins. An individual modular binding domain (such as a nanobody or a DARPin) is often not large enough by itself to fully overcome the imaging challenges for small structures. Most scaffold approaches therefore involve further components to substantially increase the overall size of the particle to be visualized by cryo-EM. In one line of attack, we have used DARPins as the modular binding domain, genetically fused by way of a continuous alpha helical connection to self-assembling protein cages, to create large symmetric scaffolds for imaging (Liu *et al*., 2018, 2019). Other studies have similarly explored the use of DARPins fused to oligomeric protein assemblies (Yao *et al*., 2019) and as structural components of larger assemblies (Vulovic *et al*., 2021).

Diverse scaffolding developments have demonstrated exciting progress, but further improvements are still needed to reliably reach high resolution for small proteins. In our previous development of protein cage-DARPin scaffolds, we reached a resolution of 3.8 Å for a 27 kDa cargo protein (Liu *et al*., 2018, 2019). In concurrent developments using antibody/nanobody approaches, the resolutions reported have been better for larger cargo proteins – e.g. 2.49 Å for a ~200 kDa receptor complex (Uchański *et al*., 2021); 2.8 Å for a 64 kDA protein (Herzik *et al*., 2019), 3.03 Å for a 58 kDa protein (Cater *et al*., 2021); ~3.2 Å for a 52 kDa protein (Fan *et al*., 2019), 3.47 Å and 3.78 Å for ~50 kDa proteins using NabFabs (Bloch *et al*., 2021). For proteins smaller than 50 kDa, the finest resolution so far is roughly 3.2 Å for a 23 kDa protein bound to a scaffold ensemble* (Wu and Rapoport, 2021).

The present study follows on our previous protein cage-DARPin scaffolding approach, retaining key advantages of those systems (*i.e*. very large size, high particle symmetry and mitigation of preferred orientation problems), while critically addressing the problem of attachment flexibility. We show that an alternative design allows for substantial rigidification of the DARPin attachments by the introduction of new protein-protein interfaces using computational sequence design. Using the key oncogenic protein KRAS, we show that the rigidification provided by this new engineering strategy, combined with underlying advantages of symmetric cage scaffolds, makes it possible to reach 3 Å resolution, sufficient to visualize point mutations and bound ligands in atomic detail. This advance demonstrates the maturation of a modular scaffold to enable routine cryo-EM structure determination of proteins even smaller than 20 kDa.

* An overall resolution of 3.2 Å was reported for the overall complex between the scaffold and KDELR (the protein of interest). A range of 3.0 to 3.5 Å was estimated for the KDELR protein itself.

## Results

### Rigidification of the imaging scaffold by interface design

Our previous cage-scaffold designs reached a resolution of 3.8 Å for the attached cargo protein (Liu *et al*., 2018, 2019), but residual flexibility in the continuous alpha helical linker fusing the cage protein subunit and the modular DARPin domain prevented higher resolution. The rigidified scaffolds we develop in the present study are based on a different cage core compared to previous publications (Liu *et al*., 2018; Liu *et al*., 2019), combined with a novel method for rigidifying the attachment of the DARPin to the cage core subunit (see Fig. 1 and Methods). We retained the idea of a continuous alpha helical connection as the primary means of rigid attachment to the symmetric cage core, and supplemented it with additional designed protein-protein interactions. In the earlier design, the individual DARPin arms – 12 in total emanating from the tetrahedrally symmetric cage – protruded independently, isolated from each other and thus permitting some residual flexibility. To make further stabilizing contacts possible, we investigated alternative design choices. A different tetrahedral protein cage known as T33-51 (Cannon *et al*., 2020), when modeled with alpha helical linkers to DARPins, oriented the protruding DARPin arms to be in near-contact with each other; three DARPins come together at each of the four vertices of the tetrahedron (Fig. 1). Then, computational interface design was used to select new amino acid sequences at the interfaces formed between three symmetry-related copies of the DARPin (see Fig. 2 and Methods). The designed interfaces between protruding DARPins confer additional stability to these key binding components of the scaffold. With this strategy, isolated flexible protrusions are eliminated, to create a much more rigid framework (Fig. 1).

**Figure 1.**
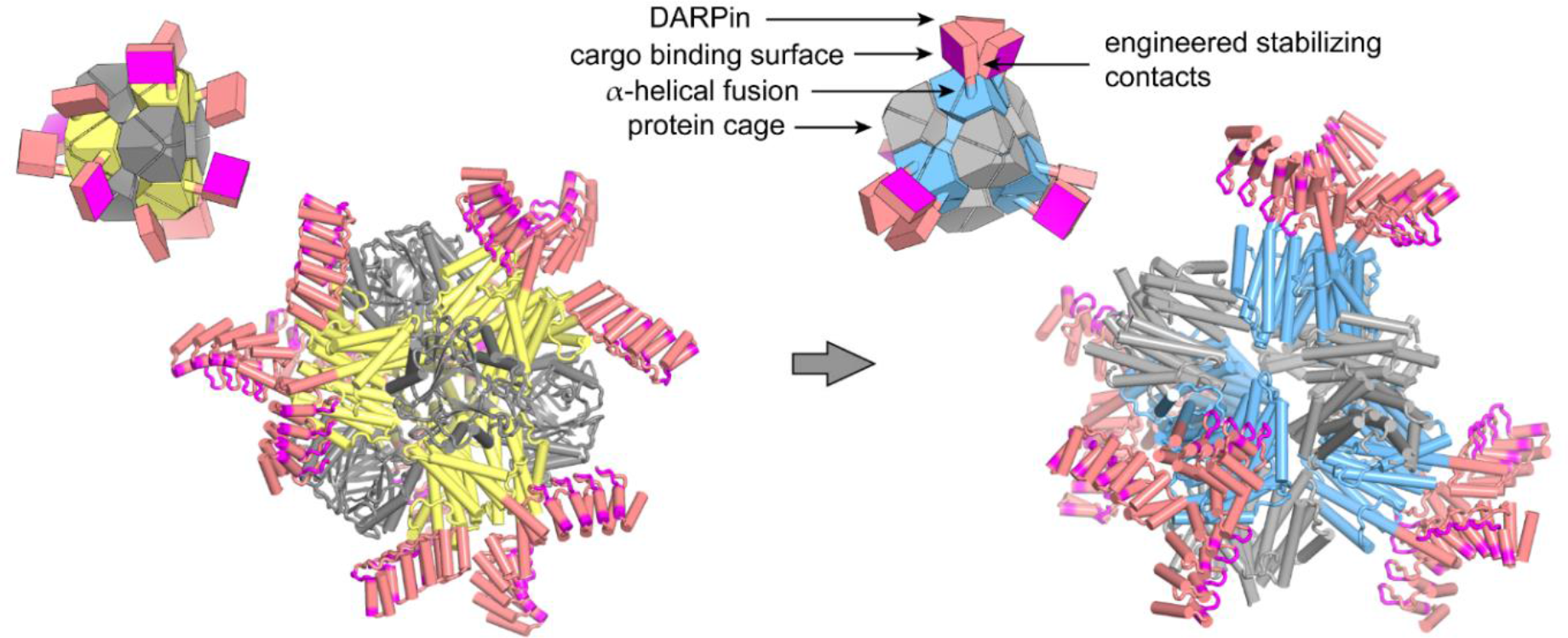
A scheme for rigidifying a modular cryo-EM imaging scaffold. (left) A previously described scaffold (Liu, *et al*. 2018; Liu *et al*. 2019), based on a self-assembling protein cage, displayed protruding DARPin domains as modular binders via continuous alpha helical fusions. The cage subunits bearing the continuous alpha helical fusion are shown in yellow. The other subunit type in this 2-component cage is shown in gray. DARPin domains are colored in salmon with their hyper variable binding regions highlighted in magenta. (right) A redesigned scaffold based on similar principles, but with protruding DARPin arms disposed to make additional protein-protein contacts with symmetric copies of each other. Designed surface mutations at the new interface away from the hypervariable region stabilize the DARPin domain, allowing high resolution cryo-EM imaging of bound cargo. The insets provide simplified geometric diagrams of the scaffold constructions.

**Figure 2.**
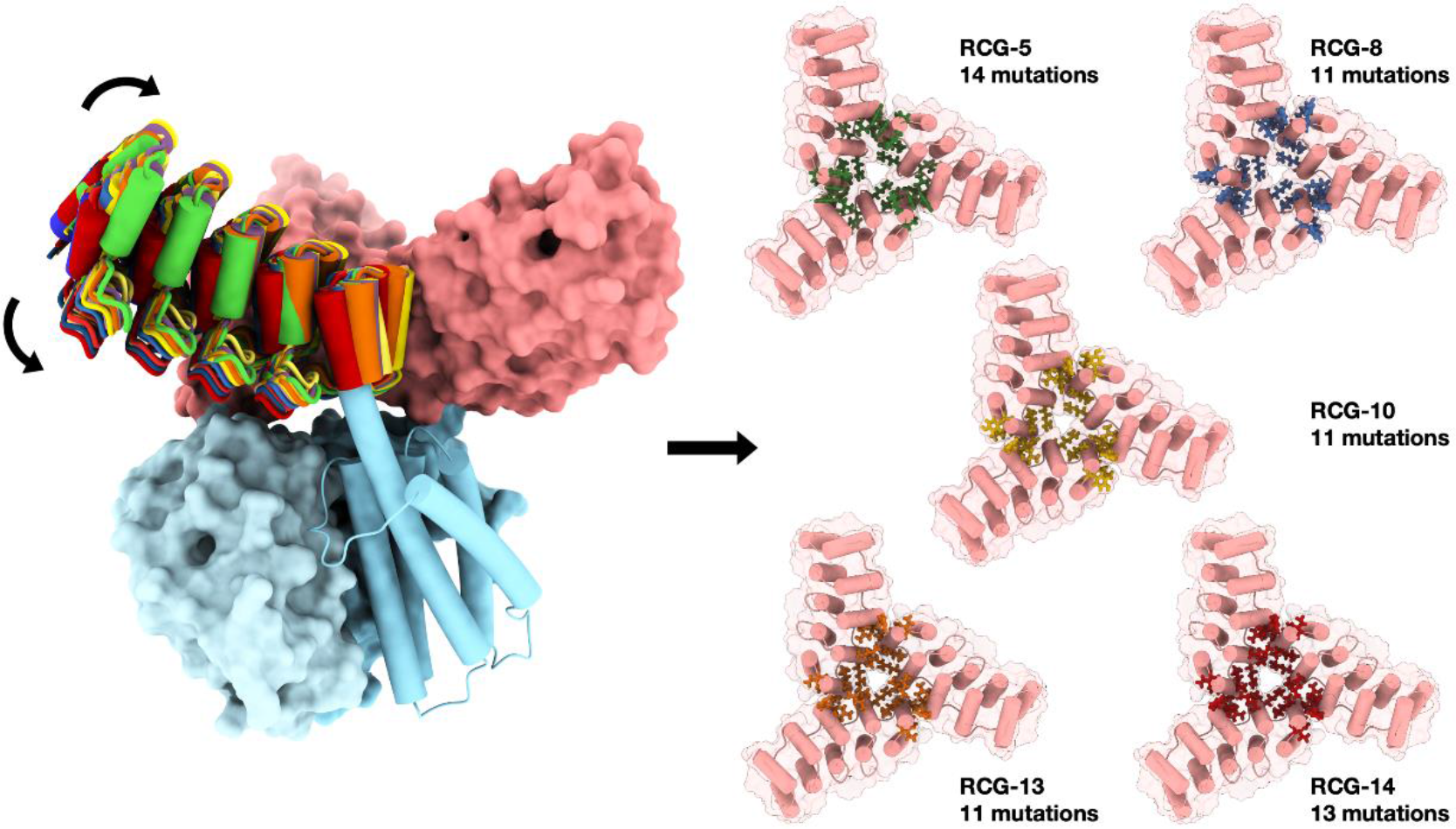
Rigidification of the designed scaffold by computational interface design between contacting DARPins. (left) Three protruding DARPins are shown fused to their protein cage subunits (blue) by a continuous alpha helical linker. A limited degree of natural flexibility (*i.e*. deviation from ideal helical parameters) was modeled in the alpha helix in order to generate distinct backbone models that sample slightly different modes of association in the new interface. One of the DARPins, subject to flexible modeling, is shown by multicolor models; the other two contacting DARPins are shown in salmon. Among top-scoring models from computational sequence design, five candidates (right) were found suitable for evaluation by cryo-EM. Mutated residues are shown in stick representation for each of the DARPins in a trimeric bundle.

Computational sequence design (see Methods) was used to generate a series of scaffold design candidates whose sequences varied in the new interface. Twelve candidates were chosen for gene synthesis and expression in *Escherichia coli* (see Methods). For comparison to earlier trials (and prior to investigating more precious cargo proteins), we chose to conduct optimization tests on scaffolds where the DARPin sequence was one that binds the model protein, super folder green fluorescent protein (sfGFP). Of the 12 designed sequences, five exhibited the two expected protein bands in SDS PAGE gels (corresponding to cage subunit A and B) after purifying presumptive protein cages by nickel affinity chromatography (Fig. S1). [Note that the cage core T33-51 is composed of subunits A and B, with stoichiometry A_12_B_12_, and cage subunit B is fused to the DARPin by way of a continuous alpha helix]. Each of these successful designs assembled as expected, according to their elution volumes in size exclusion chromatography and by negative stain TEM (Fig S2). All designs bound GFP as anticipated.

### Single-particle cryo-EM analysis of rigidified scaffolds binding GFP

To analyze the structures of the newly designed scaffolds, we collected a cryo-EM dataset for each of the five designs. Each of the cages was purified, loaded with GFP, then subjected to identical specimen preparation and data collection on a Titan Krios. For each design, particles were selected from the collected images and subjected to 3D image reconstruction, enforcing T symmetry. Analysis of the 3D reconstructions after truncation of each dataset to the same number of particles revealed different degrees of variability in the protruding DARPins. As assessed by local resolution in initial 3D reconstructions – based on similar numbers of particles and prior to extensive particle subclassification methodologies – one design, RCG-10, provided the most rigid connection between the cage core and its DARPin arm (and the bound GFP cargo). RCG-10 was therefore selected for further analysis and image data processing (Fig. 3).

**Figure 3.**
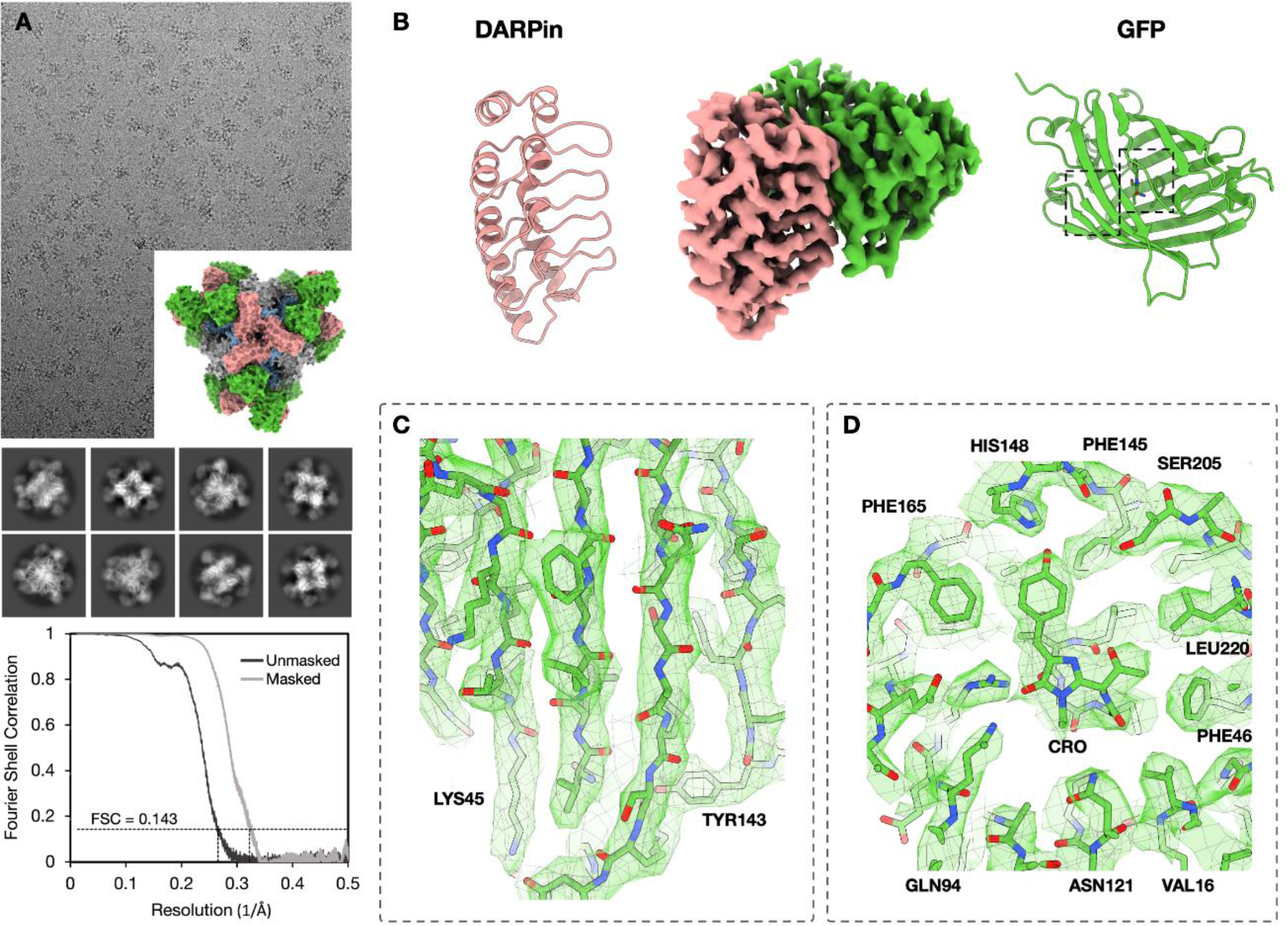
Cryo-EM structure of GFP bound to a rigidified imaging scaffold. A. Cryo-EM micrograph of the rigidified imaging scaffold bound to GFP (model shown in inset) and 2D classes from selected particles. An FSC plot illustrates agreement between independent half-maps obtained after focused classification and 3D reconstruction, masked around the GFP protein (resolution = 3.1 Å based on a correlation threshold of 0.143). B. (middle) A view of the final density map covering the DARPin and its bound GFP protein. Ribbon models of the two components are shown on the sides. C. Focused views of the density map covering several GFP beta-strands (left) and the GFP chromophore with its surrounding amino acid side chains (right).

For the imaging scaffold RCG-10 bound to GFP, ~877,000 particles were obtained from 3,575 movies. Following initial refinements applying symmetry T, the DARPin arms and attached cargo proteins were clearly visible, but still subject to some conformational variability. To visualize the bound cargo components at high resolution, we symmetry-expanded the particles (~10,000,000 particles) and performed a series of focused 3D classifications, ultimately selecting a subset of ~1,200,000 particles. Local refinements provided a reconstruction of the GFP (27 kDa) at 3.1 Å (Fig. S3 and S4). This resolution estimate is based on analysis of FSC curves (0.143 threshold) for half-maps, critically restricted to the map regions around the GFP protein. This resolution allows for facile interpretation, as illustrated by the density for the GFP chromophore and side chains from the neighboring amino acid residues (Fig. 3).

### Modular cargo binding

Given the promising results of our rigidified scaffolds bound to GFP, we explored the modularity of our new scaffold design for alternative cargoes of interest. We selected the 19.4 kDa G domain of the proto-oncogene *KRAS* as a challenging therapeutic target previously inaccessible to structure based drug discovery by cryoEM (Pantsar 2019). *KRAS* encodes a GTPase involved in signal transduction of cell proliferation pathways and is among the most frequent oncogenes in various human cancers. KRAS mutations present in approximately 25% of tumors, while driving progression in more than 60% of pancreatic and 40% of colorectal cancers (Huang *et al*., 2021). The DARPin sequence that binds KRAS, reported in prior studies (Guillard *et al*., 2017) was incorporated into the same scaffold template as above (RCG-10) to make RCG-33. For the resulting scaffold, the chimeric DARPin adopts the amino acid sequence of RCG-10 at positions making rigidifying interface contacts, while elsewhere the sequence takes its the amino acid identities from the previously established KRAS DARPin sequence. This was relatively straightforward, requiring a series of nine mutations in the backbone of the KRAS DARPin.

For imaging experiments, we investigated two different sequence variants of KRAS: single site mutants, G12V and G13C. For each, we prepared specimens containing both GDP and Mg^2+^ ligands bound to KRAS; the GDP bound form represents the inactive signaling state. To assess binding, KRAS was incubated with the designed scaffold and the resulting complex was purified by size exclusion chromatography (SEC) prior to cryo-EM analysis. The resulting chromatograms demonstrated equimolar binding between the new scaffold and KRAS. The modularity of the scaffolding system is highlighted by the demonstrated ability to mutate the variable loop sequences of the DARPin to bind diverse imaging targets (Fig. S5).

### KRAS structure analysis

The RCG-33 scaffold bound to KRAS exhibited excellent behavior under cryo-EM. For mutant G13C, ~665,000 particles were obtained from 2,000 movies. Following similar data processing as before, we obtained an overall resolution of 2.4 Å for the entire particle. We then applied symmetry expansion (7,980,000 particles) and focused 3D classification on the cargo and the DARPin and selected a subset of ~1,653,000 particles. Local refinements led to density maps where the resolution over the KRAS protein is 2.9 Å (Fig. S6 and S7). This resolution estimate is based on analysis of FSC curves (0.143 threshold) for half-maps, critically restricted to the map regions around the KRAS protein. Similarly, a density map with an estimated resolution of 3.1 Å was obtained for KRAS mutant G12V.

We evaluated the quality of the resulting maps by visual inspection and by computational means. The atomic structure of KRAS could be readily interpreted from these maps by visual analysis. Notably, density was clear for the bound GDP ligand, a Mg^2+^ ion, and surrounding amino acid residues (Fig. 4). The imaging of two different KRAS mutants allows for analysis of their atomic differences. As G12V and G13C are single site mutants, the computed difference between their density maps reveals atomic information at the two positions. Strong difference density is clear for the valine side chain at residue 12 from G12V, with difference density around the cysteine at residue 13 from G13C being somewhat weaker, possibly reflecting higher mobility (Fig. 4). We also conducted computational tests to evaluate the ability of automatic protein tracing programs (Phenix AutoBuild and Buccaneer) to build a correct structure from our density maps (Cowtan 2008, Liebschner *et al*. 2019). For the DARPin, we found that Buccaneer was able to build 97% (152 out of 156) of the backbone residues and the sequence was correctly assigned for 86% of the built residues. For KRAS G13C, main chain atoms were correctly placed for 81% (134 out of 165) and the sequence was correctly assigned over 53% of the protein. Some of the errors in automatic building of KRAS can be attributed to the program attempting to build atoms in the density originating from GDP. Subsequent manual fitting led readily to accurate and complete models for the DARPin and KRAS. Atomic differences compared to the reported X-ray structure are only 0.51 Å rmsd over the entire protein backbone. We further note that the B-factors (atomic displacement parameters) obtained by refining the KRAS structure into the cryo-EM map are well-correlated (ρ = 0.65) to those reported by x-ray crystallography, despite the disparate constraints faced by the protein under those experiments (see Fig. S8). This highlights that the resolution and map quality obtained is high enough to allow facile atomic interpretation of protein structures, as well as potentially important dynamic information.

**Figure 4.**
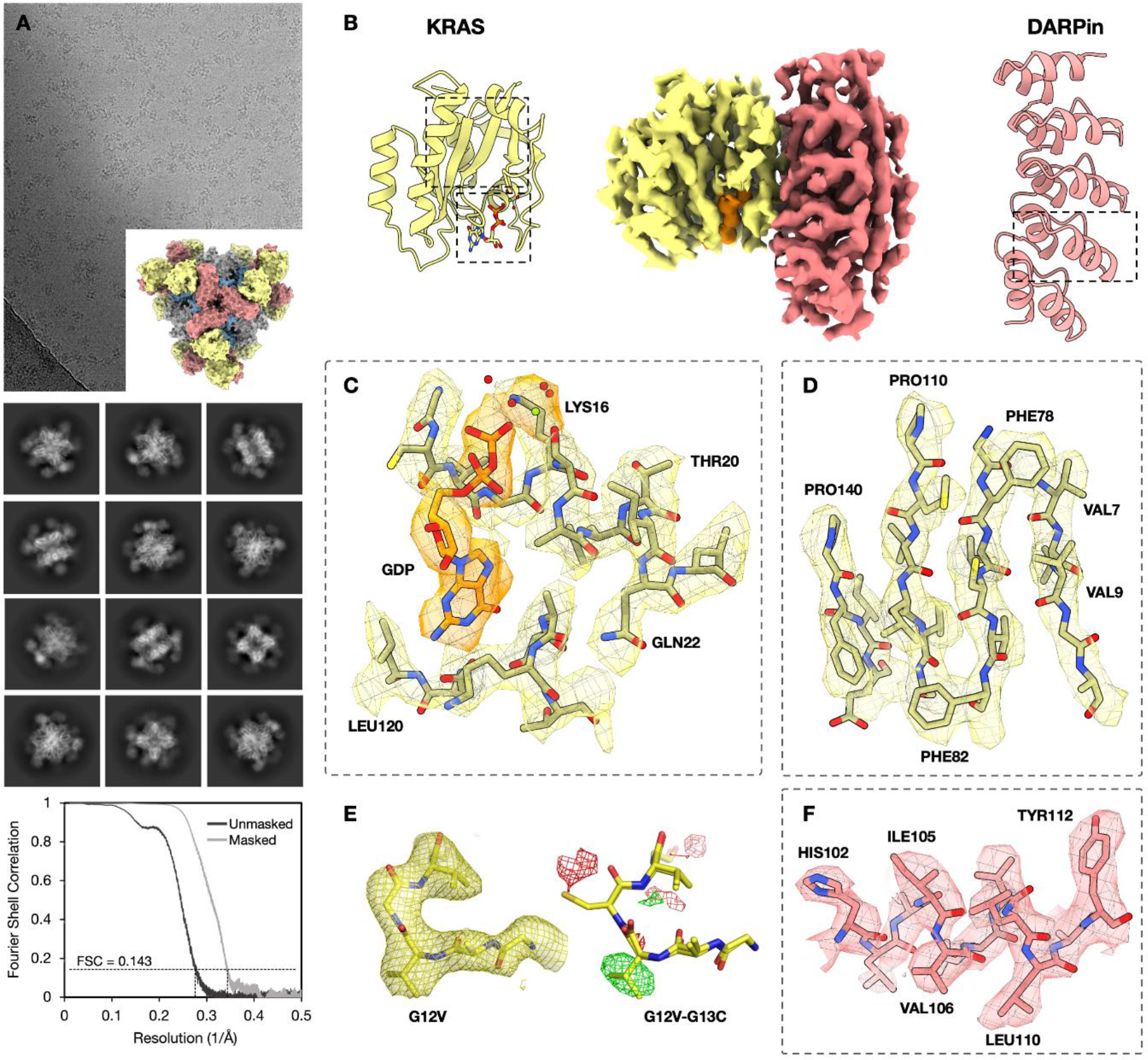
Cryo-EM structure of KRAS on a rigidified imaging scaffold. A. Cryo-EM micrograph of the rigidified imaging scaffold bound to KRAS (model shown in inset) and 2D classes from the selected particles. An FSC plot illustrates agreement between independent half-maps, obtained after focused classification and 3D reconstruction, masked around the KRAS protein (resolution = 2.9 Å based on a correlation threshold of 0.143). B. (middle) 3D reconstruction of a density map covering the DARPin and its bound KRAS protein. The GDP ligand is shown in orange. Ribbon models of the two components are shown on the sides. C, D. Focused views of the density map covering the bound GDP ligand and select regions of the KRAS structure. E. Density maps around the single site KRAS mutations analyzed. (right) A difference map (G12V minus G13C) between the two reconstructions shows strong positive features (green) corresponding to atoms only present in the KRAS G12V (*i.e*. the valine side chain) and negative features (red), corresponding to atoms only present in the KRAS G13C (*i.e*. the cysteine side chain). F. A focused view around a select region of the DARPin.

## Discussion

Our results demonstrate a new cryo-EM imaging scaffold able to reach 3 Å resolution for small proteins, even below 20 kDa. Molecular engineering, based on basic ideas for mechanical stability, combined with computational sequence design, substantially reduced the flexibility of the protruding elements of the scaffold, and thus the attached cargo proteins, enabling this gain in resolution. The current 3 Å resolution achievement provides the atomic detail required to reliably determine the structures of novel proteins. Furthermore, the approach enables routine determination of protein-ligand complex structures of challenging crystallizability, which in turn paves the road for structure based drug discovery applications for such targets. The modularity of the system – which we demonstrate here by imaging different proteins based on minor modifications to the same scaffold – makes it useful for a broad range of protein targets. The only requirement is the ability to obtain a DARPin sequence that binds the target cargo protein, which has been demonstrated in numerous prior studies (Binz *et al*., 2004, Rothenberger *et al*., 2022).

From the pursuit of diverse scaffolding approaches, various advantages and disadvantages will emerge. We anticipate that the cage-based scaffolds developed here will offer advantages related to size and symmetry. Their large size (e.g. ~900 kDa for the present scaffold when bound to KRAS) contributes to facile image processing. In addition, though experience shows that the underlying tetrahedral particle symmetry must ultimately be relaxed in order to obtain the best resolution for the attached cargo, the (near) symmetry properties of the scaffold are nonetheless valuable. A single particle presents 12 copies of the cargo, thereby tending to provide EM images with very large numbers of individual views of the cargo protein. In addition, the high symmetry of the cage means that an individual particle presents the cargo protein in 12 different orientations, strongly mitigating preferred orientation effects that challenge many cryo-EM studies. Another feature, not yet fully explored, is that each cage particle will present its bound cargo molecules at different z-heights (*i.e*. with different dispositions relative to the air-water interface). Surface forces, causing protein deformation or denaturation at the air-water interface, are believed to be a major challenge for structural integrity and ultimate image quality in cryo-EM (D’Imprima *et al*., 2019; Noble *et al*., 2018).

The versatility of the present scaffolding system opens the possibility for high throughput structural biology of small proteins. Owing to favorable properties of the present system, essentially no protein is too small. In particular, many cellular proteins of medical and therapeutic interest will be suitable for structural study with this approach. We show that here using KRAS as an exciting example of future possibilities.

## Materials and Methods

### Conformational sampling of rigidified scaffolds

The N-terminal helix of DARP14-3G124Mut5 (Liu *et al*., 2019) was spatially aligned to the C-terminal helix of each subunit from the T33-51 cage (Cannon *et al*., 2020). Using local programs, superpositions were performed between the first five helical residues of the DARPin to five residue windows from the terminal helical region of the protein cage, with different choices for the alignment segment from the protein cage. Following superposition, each conformation was evaluated for detrimental overlapping collisions and potentially favorable contacts in the fully assembled symmetric environment using local programs as well as visual inspection. Promising conformations – those where multiple protruding DARPin arms came into close proximity – were subjected to further conformational exploration by allowing for minor helix flexing. Modeling of allowable deviations from ideal alpha helix geometry was based on natural deviations observed in a large set of alpha helices extracted from high resolution crystal structures.

### Interface design calculation

All calculations were performed in the context of tetrahedral symmetry. For each sampled alignment and helical bend conformation, the resulting pose was relaxed into the REF2015 score function (Alford *et al*., 2017) using the FastRelax mover (Nivón *et al*., 2013). Then, residues in the aligned helical fusion as well as any residues located in cage subunits or other DARPins (excluding variable loop regions) within 8 Å of the aligned DARPin were marked as designable. Further, all residues within 8 Å of designable residues were designated as packable. Sequence design trajectories were performed with a coordinate constraint applied to backbone atoms using Rosetta FastDesign with the InterfaceDesign2019 protocol (Maguire *et al*., 2021) and REF2015 score function. We collected interface design metrics to quantify the resulting design success as compared to native interfaces (Janin *et al*., 2008). After analysis of the global design pool, we removed entire poses from consideration where the average design trajectory had a measured shape complementarity below 0.6, leaving eight viable poses for sampling sequence variations. Next, we ranked the design trajectories from each passing pose by applying a linear weighting scheme to the normalized metrics from each pose. These consisted of favoring fewer buried unsatisfied hydrogen bonds, lower interface energy (between complexed and unbound forms), higher interface shape complementarity, and lower interface solvation energy. Each normalized metric was equally weighted and summed to rank each trajectory. Finally, by examining the sequence diversity of the top candidates from each pose, we removed redundant sequence mutation patterns and selected 12 individual designs for characterization.

### Cloning and protein expression

The sequences of the imaging scaffolds used in this paper are described in the Supplementary Information. DNA fragments carrying the designed imaging scaffold sequences were synthesized (Integrated DNA Technologies and Twist Bioscience) and separately cloned into the vectors pET-22b (subunitB-DARPin) or pSAM (subunitA) (gifted from Jumi Shin, Addgene plasmid #45174;http://n2t.net/addgene:45174; RRID:Addgene_45174). The superfolder GFP V206A (sfGFP V206A) vector was previously described (Liu *et al*., 2019). DNA manipulations were carried out in *Escherichia coli* XL2 cells (Agilent). The proteins were expressed in *E. coli* BL21(DE3) cells (New England Biolabs) in Terrific Broth at 18°C overnight with 0.5 mM IPTG induction at an OD_600_ of 1.0.

Upon collection of the cells, pellets were resuspended in buffer (50 mM Tris, 300 mM NaCl, 20 mM imidazole, pH 8.0) supplemented with benzonase nuclease, 1 mM PMSF, EDTA-free protease inhibitor cocktail (Thermo Scientific) and 0.1% LDAO and lysed using an EmulsiFlex C3 homogenizer (Avestin). The cell lysate was cleared by centrifugation at 20,000 xg for 20 min at 4°C, the resulting supernatant was recovered and centrifuged at 10,000 xg for 10 min at 4°C and then loaded onto a HisTrap column (GE Healthcare) pre-equilibrated with the same resuspension buffer. The imaging scaffold was eluted with a linear gradient to 300 mM imidazole. Upon elution, 5 mM EDTA and 5 mM BME were added immediately for designs 5, 8, 10, 13 and 14. The eluted proteins were concentrated using Amicon Ultra-15 100 kDa molecular weight cutoff for the imaging scaffold and 3 kDa molecular weight cutoff for the GFP protein. The concentrated proteins were further purified by size exclusion chromatography using a Superose 6 Increase column, eluted with 20 mM Tris pH 8.0, 100 mM NaCl, 5 mM BME, 5 mM EDTA for designs 5, 8, 10, 13 and 14 and 20 mM Tris pH 8.0, 100 mM NaCl for design 33. Chromatography fractions were analyzed by SDS-PAGE and negative stain EM for the presence of the imaging scaffold. KRAS G12V and KRAS G13C proteins were prepared as previously described in Kettle *et al*., 2020.

### Negative stain EM

The concentration of a 3.5 μl sample of fresh Superose 6 Increase eluent was adjusted to ~100 μg/ml, applied to glow-discharged Formvar/Carbon 400 mesh Cu grids (Ted Pella Inc) for one minute and blotted to remove excess liquid. After a wash with filtered MilliQ water, the grid was stained with 2% uranyl acetate for one minute. Images were taken on a Tecnai T12, a T20, a TF20 and a Talos F200C.

### Cryo-EM data collection

Concentrated imaging scaffolds (1-10 mg/ml) were mixed with the GFP cargo to a molar ratio of 1:2 and diluted to a final concentration of 0.5-0.7 mg/ml. The final buffer composition was 20 mM Tris pH 8.0, 100 mM NaCl. Quantifoil 300 mesh R2/2 copper grids were glow discharged for 30 s at 15 mA using a PELCO easiGLow. A 3.5 μl volume of the sample was applied to the grid and then blotted and plunge-frozen into liquid nitrogen-cooled liquid ethane using a Vitrobot Mark IV (FEI). Cryo-EM data were collected on a Gatan K3 Summit direct electron detector on a Titan Krios (FEI). Images were recorded with Leginon (Suloway *et al*., 2005) and SerialEM (Mastronarde 2005) with a pixel size of 1.1 Å for designs 5, 8, 10, 13, 14, 33 (G13C) datasets and 0.856 Å for design 33 (G12V) dataset, over a defocus range of −2.5 to −0.4μm.

### Cryo-EM data processing and model building

Motion correction, CTF estimation, particle picking, 2D classification and further data processing were performed with cryoSPARC v.3.2 (Punjani *et al*., 2017). An initial set of particles was automatically picked using a blob-picker protocol. The extracted particles were 2D classified after which an *ab initio* reconstruction was generated. This reconstruction was then used for the 3D refinements enforcing T symmetry. The 3D structure was used to generate 2D projections of the particles and then used to repick the particles from the images using a template picker. The picked particles were extracted from the micrographs and went through 3D refinements enforcing T symmetry. The symmetry was then expanded, followed by further 3D classification and local refinements. For the GFP imaging scaffold we obtained an overall resolution of 2.7 Å for the entire particle and a resolution of 3.1 Å over the GFP protein, based on an FSC threshold of 0.143. For the KRAS G13C imaging scaffold we obtained an overall resolution of 2.4 Å for the entire particle and the resolution over the KRAS protein was 2.93 Å. Models were built into the maps and subjected to multiple rounds of refinement with Coot (Emsley *et al*., 2010) and PHENIX (Liebschner *et al*., 2019).

### FSC calculation

Fourier shell correlation (FSC) plots were generated using the *mtriage* tool of Phenix (Afonine *et al*. 2018). Each refined model and final map were submitted to *mtriage* along with two half maps. Masked curves correspond to the use of a smoothed mask to perform FSC calculation only around the model (Pintilie, G. *et al*. 2016).

## Acknowledgements

This work was supported by NIH grant R01GM129854 (TOY). Additional resources for sample preparation and electron microscopy screening were supported by DOE grant DE-FC02-02ER63421. Data was acquired at the Electron Imaging Center for Nanomachines (EICN) at the University of California, Los Angeles California for NanoSystems Institute (CNSI). We thank David Strugatsky and Peng Ge for assistance in cryo-EM data collection. We also thank Kevin Cannon, Ivo Atanasov, and Wong Hoi Hui for training in cryo-EM. We thank Yi Xiao Jiang, Tom Dendooven and Jack Bravo for helpful discussions about cryo-EM data processing. We thank Chris Garcia and Nathanael Caveney for discussions regarding cryo-EM scaffolding.

## References

Afonine, P. V., Klaholz, B. P., Moriarty, N. W., Poon, B. K., Sobolev, O. V., Terwilliger, T. C., Adams, P. D., & Urzhumtsev, A. (2018). New tools for the analysis and validation of cryo-EM maps and atomic models. Acta crystallographica. Section D, Structural biology, 74(Pt 9), 814–840. https://doi.org/10.1107/S2059798318009324

Alford, R., Leaver-Fay, A., Jeliazkov, J. R., O’Meara, M. J., DiMaio, F. P., Park, H., Shapovalov, M. V., Renfrew, D. P., Mulligan, V., Kappel, K., Labonte, J. W., Pacella, M., Bonneau, R., Bradley, P., Dunbrack, R. L., Das, R., Baker, D., Kuhlman, B., Kortemme, T., & Gray, J. J. (2017). The Rosetta All-Atom Energy Function for Macromolecular Modeling and Design. Journal of Chemical Theory and Computation, 13(6), 3031–3048. https://doi.org/10.1021/acs.jctc.7b00125

Binz, K. H., Amstutz, P., Kohl, A., Stumpp, M. T., Briand, C., Forrer, P., Grütter, M. G., & Plückthun, A. (2004). High-affinity binders selected from designed ankyrin repeat protein libraries. Nature Biotechnology, 22(5), nbt962. https://doi.org/10.1038/nbt962

Bloch, J. S., Mukherjee, S., Kowal, J., Filippova, E. V., Niederer, M., Pardon, E., Steyaert, J., Kossiakoff, A. A., & Locher, K. P. (2021). Development of a universal nanobody-binding Fab module for fiducial-assisted cryo-EM studies of membrane proteins. Proceedings of the National Academy of Sciences of the United States of America, 118(47), e2115435118. https://doi.org/10.1073/pnas.2115435118

Cannon, K. A., Park, R. U., Boyken, S. E., Nattermann, U., Yi, S., Baker, D., King, N. P., & Yeates, T. O. (2020). Design and structure of two new protein cages illustrate successes and ongoing challenges in protein engineering. Protein science: a publication of the Protein Society, 29(4), 919–929. https://doi.org/10.1002/pro.3802

Cater, R. J., Chua, G. L., Erramilli, S. K., Keener, J. E., Choy, B. C., Tokarz, P., Chin, C. F., Quek, D., Kloss, B., Pepe, J. G., Parisi, G., Wong, B. H., Clarke, O. B., Marty, M. T., Kossiakoff, A. A., Khelashvili, G., Silver, D. L., & Mancia, F. (2021). Structural basis of omega-3 fatty acid transport across the blood-brain barrier. Nature, 595(7866), 315–319. https://doi.org/10.1038/s41586-021-03650-9

Coscia, F., Estrozi, L. F., Hans, F., Malet, H., Noirclerc-Savoye, M., Schoehn, G., & Petosa, C. (2016). Fusion to a homo-oligomeric scaffold allows cryo-EM analysis of a small protein. Scientific reports, 6, 30909. https://doi.org/10.1038/srep30909.

Cowtan K. (2008). Fitting molecular fragments into electron density. Acta crystallographica. Section D, Biological crystallography, 64 (Pt 1), 83–89. https://doi.org/10.1107/S0907444907033938

D’Imprima, E., Floris, D., Joppe, M., Sánchez, R., Grininger, M., & Kühlbrandt, W. (2019). Protein denaturation at the air-water interface and how to prevent it. eLife, 8, e42747. https://doi.org/10.7554/eLife.42747

Emsley, P., Lohkamp, B., Scott, W. G., & Cowtan, K. (2010). Features and development of Coot. Acta crystallographica. Section D, Biological crystallography, 66(Pt 4), 486–501. https://doi.org/10.1107/S0907444910007493

Fan, X., Wang, J., Zhang, X., Yang, Z., Zhang, J. C., Zhao, L., Peng, H. L., Lei, J., & Wang, H. W. (2019). Single particle cryo-EM reconstruction of 52 kDa streptavidin at 3.2 Angstrom resolution. Nature communications, 10(1), 2386. https://doi.org/10.1038/s41467-019-10368-w

Guillard, S., Kolasinska-Zwierz, P., Debreczeni, J., Breed, J., Zhang, J., Bery, N., Marwood, R., Tart, J., Overman, R., Stocki, P., Mistry, B., Phillips, C., Rabbitts, T., Jackson, R., & Minter, R. (2017). Structural and functional characterization of a DARPin which inhibits Ras nucleotide exchange. Nature communications, 8, 16111. https://doi.org/10.1038/ncomms16111

Henderson R. (1995). The potential and limitations of neutrons, electrons and X-rays for atomic resolution microscopy of unstained biological molecules. Quarterly reviews of biophysics, 28(2), 171–193. https://doi.org/10.1017/s003358350000305x

Herzik, M. A., Jr, Wu, M., & Lander, G. C. (2019). High-resolution structure determination of sub-100 kDa complexes using conventional cryo-EM. Nature communications, 10(1), 1032. https://doi.org/10.1038/s41467-019-08991-8

Huang, L., Guo, Z., Wang, F., & Fu, L. (2021). KRAS mutation: from undruggable to druggable in cancer. Signal transduction and targeted therapy, 6(1), 386. https://doi.org/10.1038/s41392-021-00780-4

Janin, J., Bahadur, R. P., & Chakrabarti, P. (2008). Protein–protein interaction and quaternary structure. Quarterly Reviews of Biophysics, 41(2), 133–180. https://doi.org/10.1017/s0033583508004708

Kettle, J. G., Bagal, S. K., Bickerton, S., Bodnarchuk, M. S., Breed, J., Carbajo, R. J., Cassar, D. J., Chakraborty, A., Cosulich, S., Cumming, I., Davies, M., Eatherton, A., Evans, L., Feron, L., Fillery, S., Gleave, E. S., Goldberg, F. W., Harlfinger, S., Hanson, L., Howard, M., … Zhao, X. (2020). Structure-Based Design and Pharmacokinetic Optimization of Covalent Allosteric Inhibitors of the Mutant GTPase KRAS^G12C^. Journal of medicinal chemistry, 63(9), 4468–4483. https://doi.org/10.1021/acs.jmedchem.9b01720

Kratz, P. A., Böttcher, B., & Nassal, M. (1999). Native display of complete foreign protein domains on the surface of hepatitis B virus capsids. Proceedings of the National Academy of Sciences of the United States of America, 96(5), 1915–1920. https://doi.org/10.1073/pnas.96.5.1915.

Liebschner, D., Afonine, P. V., Baker, M. L., Bunkóczi, G., Chen, V. B., Croll, T. I., Hintze, B., Hung, L. W., Jain, S., McCoy, A. J., Moriarty, N. W., Oeffner, R. D., Poon, B. K., Prisant, M. G., Read, R. J., Richardson, J. S., Richardson, D. C., Sammito, M. D., Sobolev, O. V., Stockwell, D. H., … Adams, P. D. (2019). Macromolecular structure determination using X-rays, neutrons and electrons: recent developments in Phenix. Acta crystallographica. Section D, Structural biology, 75(Pt 10), 861–877. https://doi.org/10.1107/S2059798319011471

Liu, Y., Gonen, S., Gonen, T., & Yeates, T. O. (2018). Near-atomic cryo-EM imaging of a small protein displayed on a designed scaffolding system. Proceedings of the National Academy of Sciences of the United States of America, 115(13), 3362–3367. https://doi.org/10.1073/pnas.1718825115

Liu, Y., Huynh, D. T., & Yeates, T. O. (2019). A 3.8 Å resolution cryo-EM structure of a small protein bound to an imaging scaffold. Nature communications, 10(1), 1864. https://doi.org/10.1038/s41467-019-09836-0

Maguire, J. B., Haddox, H. K., Strickland, D., Halabiya, S. F., Coventry, B., Griffin, J. R., Pulavarti, S. V. S. R. K., Cummins, M., Thieker, D. F., Klavins, E., Szyperski, T., DiMaio, F., Baker, D., & Kuhlman, B. (2021). Perturbing the energy landscape for improved packing during computational protein design. Proteins: Structure, Function, and Bioinformatics, 4. https://doi.org/10.1002/prot.26030

Martin, T. G., Bharat, T. A., Joerger, A. C., Bai, X. C., Praetorius, F., Fersht, A. R., Dietz, H., & Scheres, S. H. (2016). Design of a molecular support for cryo-EM structure determination. Proceedings of the National Academy of Sciences of the United States of America, 113(47), E7456–E7463. https://doi.org/10.1073/pnas.1612720113

Mastronarde D. N. (2005). Automated electron microscope tomography using robust prediction of specimen movements. Journal of structural biology, 152(1), 36–51. https://doi.org/10.1016/j.jsb.2005.07.007

McMahon, C., Baier, A. S., Pascolutti, R., Wegrecki, M., Zheng, S., Ong, J. X., Erlandson, S. C., Hilger, D., Rasmussen, S., Ring, A. M., Manglik, A., & Kruse, A. C. (2018). Yeast surface display platform for rapid discovery of conformationally selective nanobodies. Nature structural & molecular biology, 25(3), 289–296. https://doi.org/10.1038/s41594-018-0028-6

Morrison, M. S., Wang, T., Raguram, A., Hemez, C., & Liu, D. R. (2021). Disulfide-compatible phage-assisted continuous evolution in the periplasmic space. Nature Communications, 12(1), 5959. https://doi.org/10.1038/s41467-021-26279-8

Nivón, L. G., Moretti, R., & Baker, D. (2013). A Pareto-optimal refinement method for protein design scaffolds. PloS one, 8(4), e59004. https://doi.org/10.1371/journal.pone.0059004

Noble, A. J., Dandey, V. P., Wei, H., Brasch, J., Chase, J., Acharya, P., Tan, Y. Z., Zhang, Z., Kim, L. Y., Scapin, G., Rapp, M., Eng, E. T., Rice, W. J., Cheng, A., Negro, C. J., Shapiro, L., Kwong, P. D., Jeruzalmi, D., des Georges, A., Potter, C. S., … Carragher, B. (2018). Routine single particle CryoEM sample and grid characterization by tomography. eLife, 7, e34257. https://doi.org/10.7554/eLife.34257

Pantsar T. (2019). The current understanding of KRAS protein structure and dynamics. Computational and structural biotechnology journal, 18, 189–198. https://doi.org/10.1016/j.csbj.2019.12.004

Pintilie, G., Chen, D.-H., Haase-Pettingell, C. A., King, J. A., & Chiu, W. (2016). Resolution and Probabilistic Models of Components in CryoEM Maps of Mature P22 Bacteriophage. Biophysical Journal, 110(4), 827–839. https://doi.org/10.1016/j.bpj.2015.11.3522

Punjani, A., Rubinstein, J. L., Fleet, D. J., & Brubaker, M. A. (2017). cryoSPARC: algorithms for rapid unsupervised cryo-EM structure determination. Nature methods, 14(3), 290–296. https://doi.org/10.1038/nmeth.4169

Rothenberger, S., Hurdiss, D. L., Walser, M., Malvezzi, F., Mayor, J., Ryter, S., Moreno, H., Liechti, N., Bosshart, A., Iss, C., Calabro, V., Cornelius, A., Hospodarsch, T., Neculcea, A., Looser, T., Schlegel, A., Fontaine, S., Villemagne, D., Paladino, M., Schiegg, D., … Trimpert, J. (2022). The trispecific DARPin ensovibep inhibits diverse SARS-CoV-2 variants. Nature biotechnology, 10.1038/s41587-022-01382-3. Advance online publication. https://doi.org/10.1038/s41587-022-01382-3

Sathiamoorthy, S., & Shin, J. A. (2012). Boundaries of the origin of replication: creation of a pET-28a-derived vector with p15A copy control allowing compatible coexistence with pET vectors. PloS one, 7(10), e47259. https://doi.org/10.1371/journal.pone.0047259

Suloway, C., Pulokas, J., Fellmann, D., Cheng, A., Guerra, F., Quispe, J., Stagg, S., Potter, C. S., & Carragher, B. (2005). Automated molecular microscopy: the new Leginon system. Journal of structural biology, 151(1), 41–60. https://doi.org/10.1016/j.jsb.2005.03.010

Uchański, T., Masiulis, S., Fischer, B., Kalichuk, V., López-Sánchez, U., Zarkadas, E., Weckener, M., Sente, A., Ward, P., Wohlkönig, A., Zögg, T., Remaut, H., Naismith, J. H., Nury, H., Vranken, W., Aricescu, A. R., Pardon, E., & Steyaert, J. (2021). Megabodies expand the nanobody toolkit for protein structure determination by single-particle cryo-EM. Nature methods, 18(1), 60–68. https://doi.org/10.1038/s41592-020-01001-6.

Vulovic, I., Yao, Q., Park, Y. J., Courbet, A., Norris, A., Busch, F., Sahasrabuddhe, A., Merten, H., Sahtoe, D. D., Ueda, G., Fallas, J. A., Weaver, S. J., Hsia, Y., Langan, R. A., Plückthun, A., Wysocki, V. H., Veesler, D., Jensen, G. J., & Baker, D. (2021). Generation of ordered protein assemblies using rigid three-body fusion. Proceedings of the National Academy of Sciences of the United States of America, 118(23), e2015037118. https://doi.org/10.1073/pnas.2015037118

Wu, X., & Rapoport, T. A. (2021). Cryo-EM structure determination of small proteins by nanobody-binding scaffolds (Legobodies). Proceedings of the National Academy of Sciences of the United States of America, 118(41), e2115001118. https://doi.org/10.1073/pnas.2115001118

Yao, Q., Weaver, S. J., Mock, J. Y., & Jensen, G. J. (2019). Fusion of DARPin to Aldolase Enables Visualization of Small Protein by Cryo-EM. Structure (London, England : 1993), 27(7), 1148–1155.e3. https://doi.org/10.1016/j.str.2019.04.003

Yeates, T. O., Agdanowski, M. P., & Liu, Y. (2020). Development of imaging scaffolds for cryo-electron microscopy. Current Opinion in Structural Biology, 60, 142–149. https://doi.org/10.1016/j.sbi.2020.01.012

