## Supplementary Materials for "Rigidified Scaffolds for 3 Angstrom Resolution Cryo-EM of Small Therapeutic Protein Targets"

Supplementary Figures:

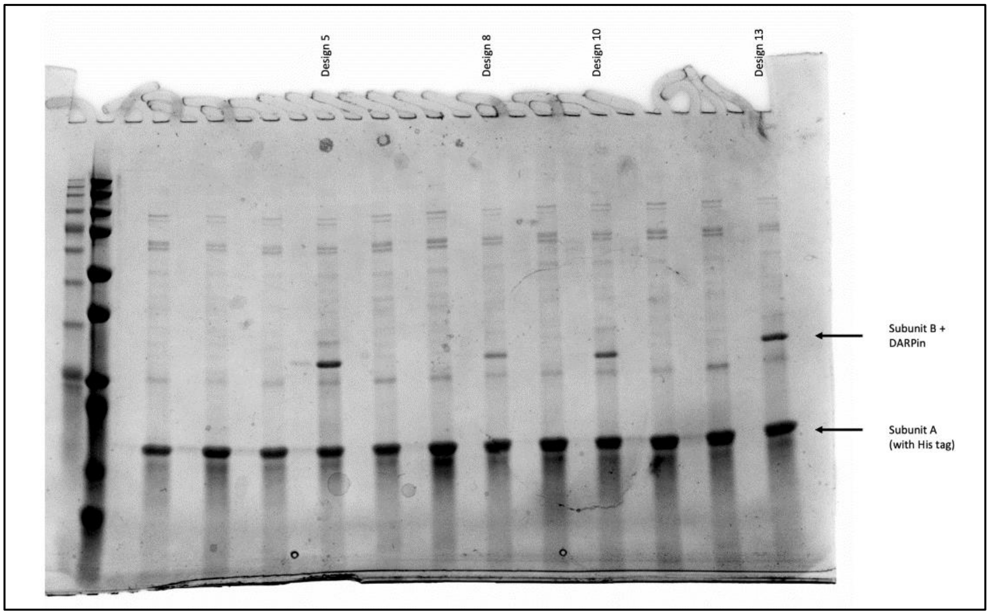


Figure S1. SDS PAGE gel showing co-elution of the two protein chains, A and B, comprising a scaffold. Subunit A (a component of the cage core) is His-tagged. Subunit B is a fusion between a cage core component and a DARPin that serves to bind diverse cargo proteins for imaging.

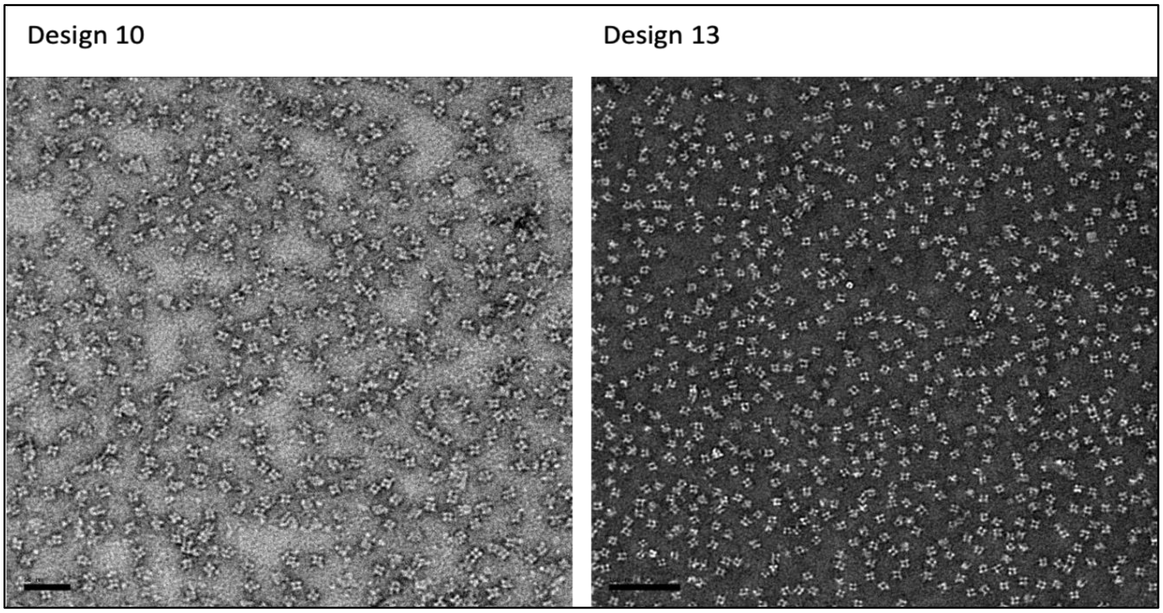


Figure S2. Negative staining of the rigidified imaging scaffold particles.

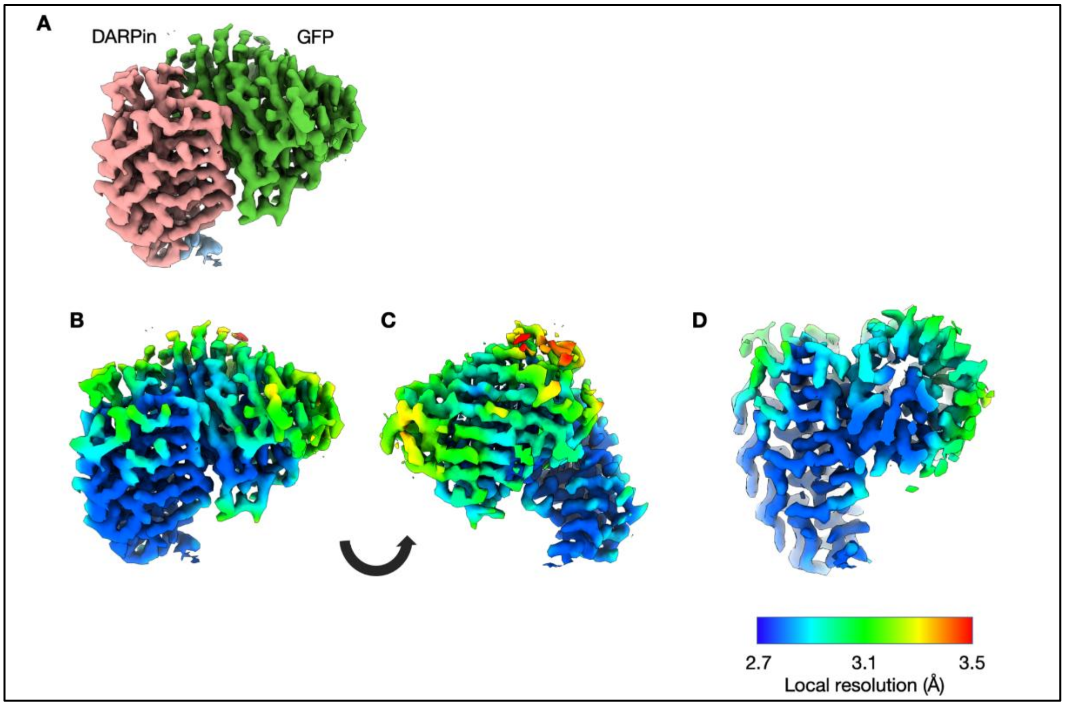


Figure S3. Local resolution for the focus-refined map of the GFP and DARPin. A. DARPin (salmon) and GFP (green). B and C, two rotated views of the map colored by local resolution. D, a cross section of the map.

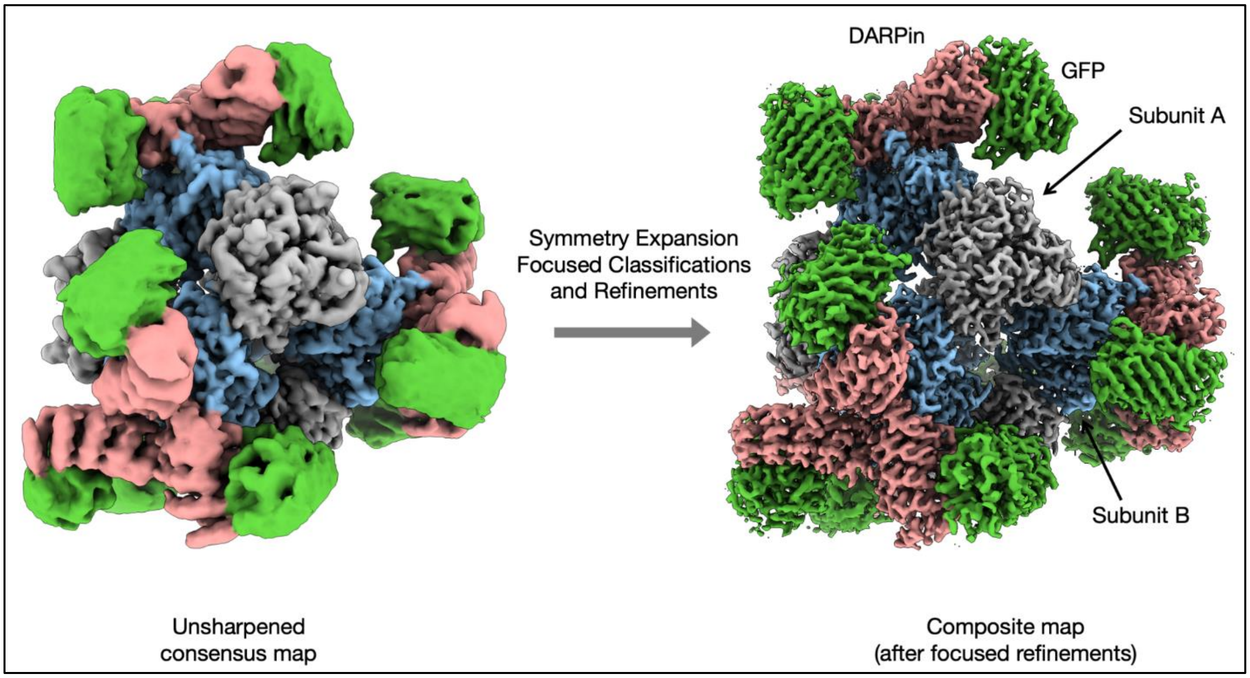


Figure S4. Cryo-EM densities for the imaging scaffold bound to GFP. The overall reconstruction (left) and a composite map of the focused refinements (right). After symmetry expansion (symmetry T), focused classifications and refinements were performed with a mask encompassing mainly one GFP (green) and one DARPin (salmon).

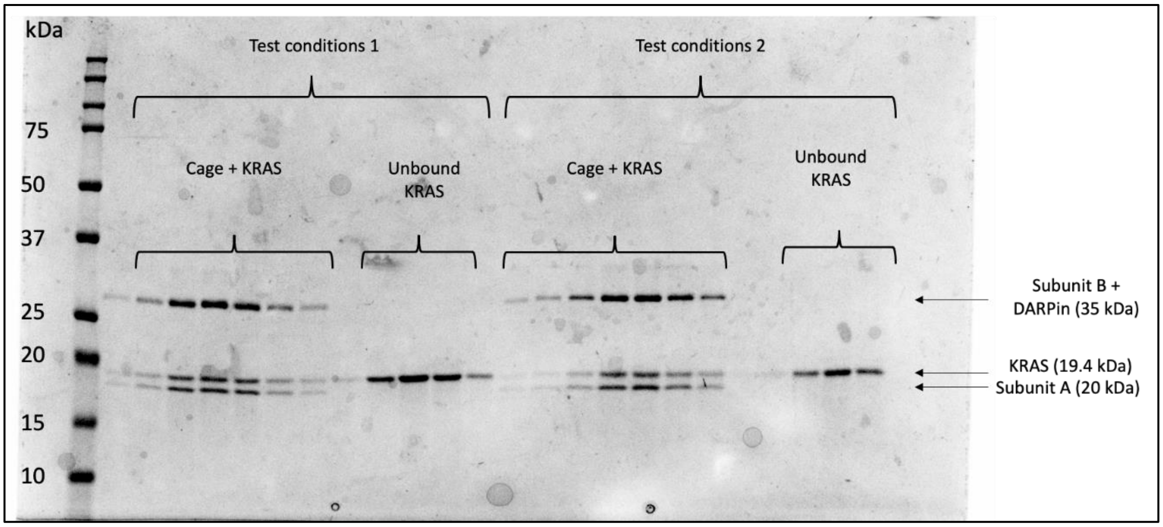


Figure S5. Binding assay of KRAS G13C to rigidified imaging scaffolds. The early SEC fractions show co-elution of the cage components with the KRAS (bound). The later fractions correspond to unbound KRAS. The higher band (~35 kDa) corresponds to the cage subunit B fused to the DARPin, the middle band corresponds to the KRAS protein (19.4 kDa) and the lower band is the cage subunit A (20kDa). Cage subunit A runs slightly smaller than its known size.

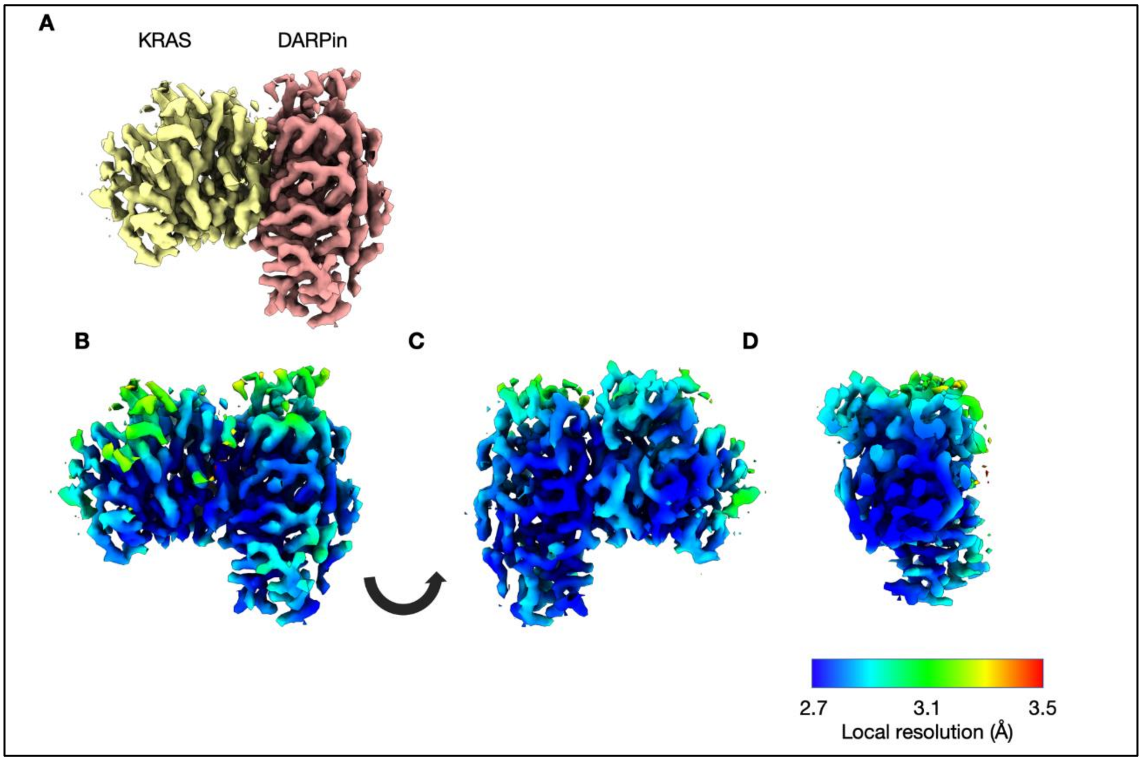


Figure S6. Local resolution for the focus-refined map of the KRAS G13C and DARPin. A. DARPin (salmon) and KRAS (yellow). B and C, two rotated views of the map colored by local resolution. D, cross section of the map.

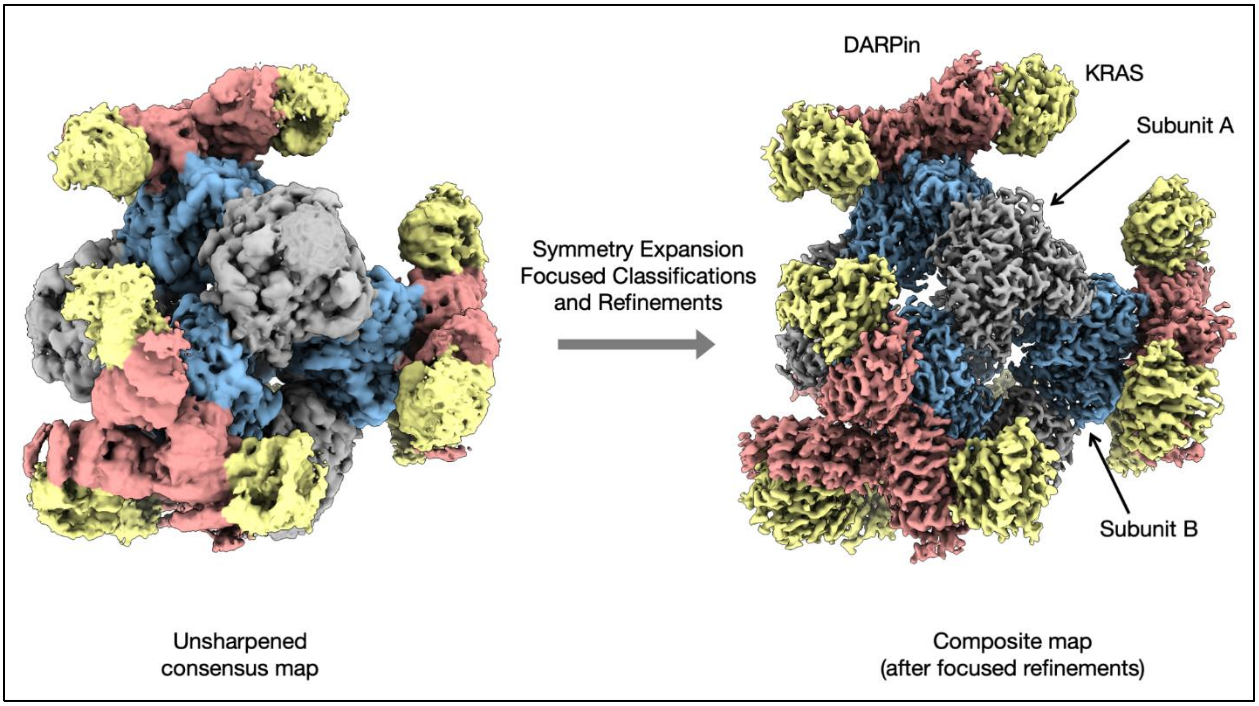


Figure S7. Cryo-EM densities for the imaging scaffold bound to KRAS G13C. The overall reconstruction (left) and a composite map of the focused refinements (right). After symmetry expansion (symmetry T), focused classifications and refinements were performed with a mask encompassing mainly one KRAS (yellow) and one DARPin (salmon).

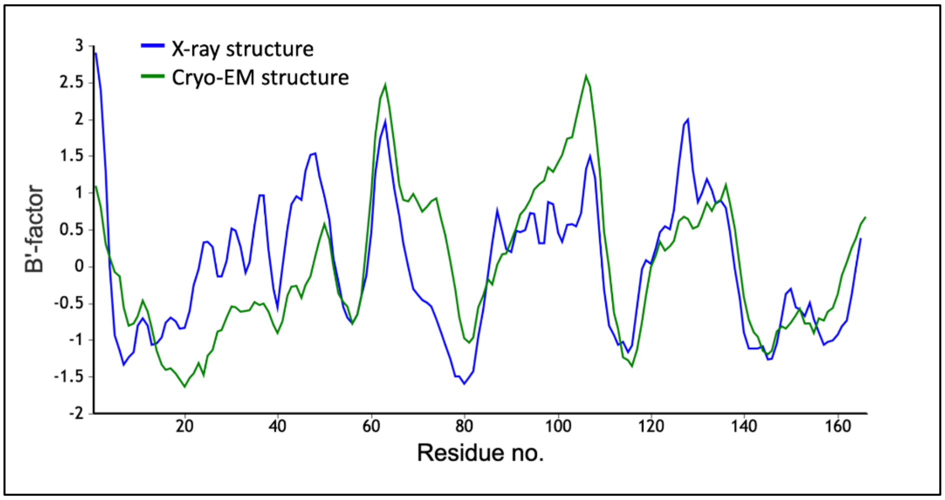


Figure S8. Plot of refined B-factors for the KRAS structure. Strong agreement is evident between the X-ray crystal structure (pdb 5o2s) and our new structure, which was built *de novo* and refined (setting B-factors to a uniform starting value of 20 Å$^2$) into the final cryo-EM map. The B-factors are averaged over individual amino acid residues and smoothed over a 3-residue window, then normalized for direct comparison using the BANΔIT toolkit (Barthels et al., 2021). The calculated correlation coefficient is 0.65.

Supplementary Notes:

**Protein sequences**

Cage Protein T33-51, Subunit A:
MRITTKVGDKGSTRLFGGEEVWKDDPIIEANGTLDELTSFIGEAKHYVDEEMKGILEEIQ
NDIYKIMGEIGSKGKIEGISEERIKWLAGLIERYSEMVNKLSFVLPGGTLESAKLDVCRTI
ARRAERKVATVLREFGIGTLAAIYLALLSRLLFLLARVIEIEKNKLKEVRS

RCG-5, Subunit B, which binds GFP:
MFTRRGDQGETDLANRARVGKDSPVVEVQGTIDELNSFIGYALVLSRWDDIRNDLFRIQ
NDLFVLGEDVSTGGKGRTVTMDMIIYLIKRSVEMKAEIGKIELFVVPGGSVESASLHMA
RAVSRRLERRIKAASELTEINANVLLYANMLSNILFMHALISNKRKEELDKKLLEAARAG
DKYAVDALLAKGADVNAADDVGVTPLHLAAQRGHLEIVEVLLKRGWDINAADLWGQ
TPLHLAATAGHLEIVELLLWYGADVNARDNIGHTPLHLAAWAGHLEIVEVLLKYGADV
NAQDKFGKTPFDLAIDNGNEDIAEVLQKAA

RCG-8, Subunit B, which binds GFP:
MFTRRGDQGETDLANRARVGKDSPVVEVQGTIDELNSFIGYALVLSRWDDIRNDLFRIQ
NDLFVLGEDVSTGGKGRTVTMDMIIYLIKRSVEMKAEIGKIELFVVPGGSVESASLHMA
RAVSRRLERRIKAASELTEINANVLLYANMLSNILFMHALISNKRKEELDKKLLEAARAG
YDDKVAWLLALGADVNAADDVGVTPLHLAAQRGHLEIVEVLLKRGADINAADLWGQ
TPLHLAATAGHLEIVEKLLRCGADVNARDNIGHTPLHLAAWAGHLEIVEVLLKYGADV
NAQDKFGKTPFDLAIDNGNEDIAEVLQKAA

RCG-10, Subunit B, which binds GFP:
MFTRRGDQGETDLANRARVGKDSPVVEVQGTIDELNSFIGYALVLSRWDDIRNDLFRIQ
NDLFVLGEDVSTGGKGRTVTMDMIIYLIKRSVEMKAEIGKIELFVVPGGSVESASLHMA
RAVSRRLERRIKAASELTEINANVLLYANMLSNILFMHALISNKRKEELDKKLLEAARAG
YDDQVAALLAKGADVNAADDVGVTPLHLAAQRGHLEIVEVLLKRGADINAADLWGQT
PLHLAATAGHLEIVELLLRWGADVNARDNIGHTPLHLAAWAGHLEIVEVLLKYGADVN
AQDKFGKTPFDLAIDNGNEDIAEVLQKAA

RCG-13, Subunit B, which binds GFP:
MFTRRGDQGETDLANRARVGKDSPVVEVQGTIDELNSFIGYALVLSRWDDIRNDLFRIQ
NDLFVLGEDVSTGGKGRTVTMDMIIYLIKRSVEMKAEIGKIELFVVPGGSVESASLHMA
RAVSRRLERRIKAASELTEINANVLLYANMLSNILFMHALISNKRKDELDKKLLEAARA
GIDDAVAALLAKGADVNAADDVGVTPLHLAAQRGHLEIVKVLLLRGADINAADLWGQ
TPLHLAATAGHLEIVELLLRCGADVNARDNIGHTPLHLAAWAGHLEIVEVLLKYGADV
NAQDKFGKTPFDLAIDNGNEDIAEVLQKAA

RCG-14, Subunit B, which binds GFP:
MFTRRGDQGETDLANRARVGKDSPVVEVQGTIDELNSFIGYALVLSRWDDIRNDLFRIQ
NDLFVLGEDVSTGGKGRTVTMDMIIYLIKRSVEMKAEIGKIELFVVPGGSVESASLHMA
RAVSRRLERRIKAASELTEINANVLLYANMLSNILFMHALISNKRRDERNKKLLEAARA
GIDDAVDWLLALGADVNAADDVGVTPLHLAAQRGHLEIVKVLLSRGADINAADLWGQ
TPLHLAATAGHLEIVELLLRCGADVNARDNIGHTPLHLAAWAGHLEIVEVLLKYGADV
NAQDKFGKTPFDLAIDNGNEDIAEVLQKAA

RCG-33, Subunit B, which binds KRAS:
MFTRRGDQGETDLANRARVGKDSPVVEVQGTIDELNSFIGYALVLSRWDDIRNDLFRIQ
NDLFVLGEDVSTGGKGRTVTMDMIIYLIKRSVEMKAEIGKIELFVVPGGSVESASLHMA
RAVSRRLERRIKAASELTEINANVLLYANMLSNILFMHALISNKRKEELDKKLLEAARAG
QDDEVAALLAKGADVNAHDTFGFTPLHLAALYGHLEIVEVLLKRGADINADDSYGRTP
LHLAAMRGHLEIVELLLRWGADVNAADEEGRTPLHLAAKRGHLEIVEVLLKNGADVN
AQDKFGKTAFDISIDNGNEDLAEILQKL

Supplementary References:

Barthels, F., Schirmeister, T., & Kersten, C. (2021). BANΔIT: B'-Factor Analysis for Drug Design and Structural Biology. Molecular informatics, 40(1), e2000144. https://doi.org/10.1002/minf.202000144
